# Small RNAs gained during epididymal transit of sperm are essential for embryonic development in mice

**DOI:** 10.1101/311670

**Authors:** Colin C. Conine, Fengyun Sun, Lina Song, Jaime A. Rivera-Pérez, Oliver J. Rando

## Abstract

The small RNA payload of mammalian sperm undergoes dramatic remodeling during development, as several waves of microRNAs and tRNA fragments are shipped to sperm during post-testicular maturation in the epididymis. Here, we take advantage of this developmental process to probe the function of the sperm RNA payload in preimplantation development. We generated zygotes via intracytoplasmic sperm injection (ICSI) using sperm obtained from the proximal (caput) vs. distal (cauda) epididymis, then characterized development of the resulting embryos. Embryos generated using caput sperm significantly overexpress multiple regulatory factors throughout preimplantation development, and subsequently implant inefficiently and fail soon after implantation. Remarkably, microinjection of purified cauda-specific small RNAs into caput-derived embryos not only completely rescued preimplantation molecular defects, but also suppressed the postimplantation embryonic lethality phenotype. These findings reveal an essential role for small RNA remodeling during post-testicular maturation of mammalian sperm, and identify a specific preimplantation gene expression program responsive to sperm-delivered microRNAs.

## INTRODUCTION

In sexually-reproducing organisms, the vast majority of inheritance is mediated by the two half-complements of genetic material carried by the gametes. However, beyond the DNA blueprint provided by the gametes, other factors delivered during fertilization are required to initiate embryogenesis, and to orchestrate a robust cascade of differentiation events required for multicellular development. Many maternally supplied factors have been demonstrated to be essential for development, as for example 1) mitochondria are strictly maternally-inherited in the majority of species, 2) many maternally-supplied proteins and RNAs are required to initiate the first cellular division after fertilization, and 3) defects in imprinted gene expression underly the failure of embryos with various paternal disomies in mammals.

In contrast, due to the discrepancy in size of sperm and oocyte (one thousandfold or more in most organisms), it is generally believed that sperm contribute little or nothing to early development beyond their haploid genome, a small number of epigenetic marks that are not “erased” by maternal factors, and, in some organisms, the first functional centriole. That said, it is increasingly clear that the male gamete carries additional information that can affect offspring development (Rando 2012), and a growing number of studies in mammals have implicated sperm RNAs in control of early embryonic development and offspring phenotype. For example, embryos generated using *Dicer* cKO sperm exhibit defects in progression to the blastocyst stage of development, and these defects can be rescued by microinjection of purified sperm RNAs (Yuan et al. 2016). Moreover, we and others have found that paternal environmental conditions can alter the RNA payload of mature sperm, and in some cases injection of sperm RNAs has been shown to recapitulate either regulatory or physiological phenotypes induced in offspring of natural matings (Rassoulzadegan et al. 2006; Fullston et al. 2013; Rodgers et al. 2013; Gapp et al. 2014; Rodgers et al. 2015; Chen et al. 2016; Sharma et al. 2016). Nonetheless, although small RNAs play key roles in well-characterized transgenerational epigenetic inheritance systems such as paramutation in maize and RNA interference in plants and worms (Fire et al. 1998; Hamilton and Baulcombe 1999; Volpe et al. 2002; Zilberman et al. 2003; Arteaga-Vazquez and Chandler 2010), it remains unclear what functions are served by the small RNA payload of mammalian sperm upon delivery to the zygote.

Developing mammalian sperm exhibit multiple rounds of small RNA dynamics, starting with two major waves of piRNA production during spermatogenesis in the testis (Li et al. 2013), followed by a global loss of piRNAs and gain of 5’ tRNA fragments (tRFs) that occurs during post-testicular maturation of sperm in the epididymis (Peng et al. 2012; Garcia-Lopez et al. 2014; Chen et al. 2016; Sharma et al. 2016). Mature sperm from the cauda (distal) epididymis, which exhibit progressive motility and are competent for in vitro fertilization, carry an RNA repertoire comprised chiefly of 5’ tRFs along with a smaller population (~10% of small RNAs) of microRNAs. In addition to this dramatic switch between the dominant small RNA species that occurs as sperm enter the epididymis, the abundance of specific small RNAs varies even between sperm obtained from the proximal (caput) vs. distal (cauda) epididymis (**Figure 1A, Supplemental Figure 1**) – while caput sperm already carry high levels of tRF-Glu-CTC and tRF-Gly-GCC, levels of the similarly-abundant tRF-Val-CAC are dramatically (>10-fold) higher in cauda sperm than in caput sperm (Sharma et al. 2016). Levels of specific microRNAs also vary significantly during epididymal maturation of sperm, as for example microRNAs encoded in several genomic clusters, such as the X-linked miR-880 cluster and the miR17-92 oncomiR cluster, are far more abundant in cauda sperm than in caput sperm and appear to be delivered to maturing sperm by fusion with epididymosomes (Nixon et al. 2015; Sharma et al. 2016) (**Supplemental Figure S1B**).

**Figure 1.**
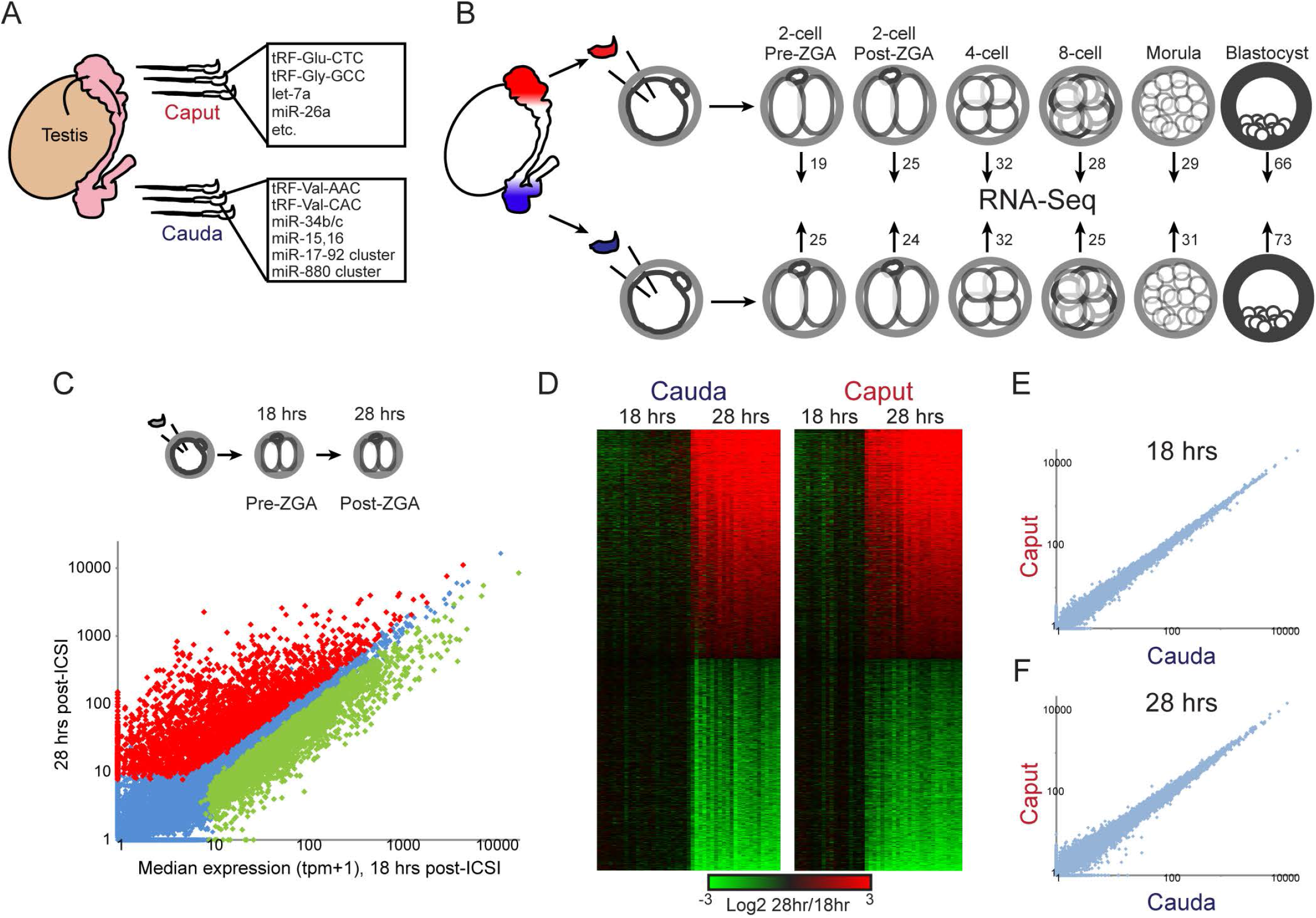
Experimental pipeline. A) Sperm obtained from the proximal (caput) and distal (cauda) epididymis carry distinct small RNA populations (Nixon et al. 2015; Sharma et al. 2016). These sperm samples differ in the abundance of hundreds of small RNAs – shown here are prominent examples of relevant RNA species. The RNAs listed for caput sperm are present at high abundance in caput sperm and are generally maintained at high levels in cauda sperm, while those listed for cauda sperm are examples of small RNAs that are scarce in caput sperm but specifically gained in cauda sperm. See also **Supplemental Figure S1**. B) Intracytoplasmic sperm injection (ICSI) pipeline. Paired caput and cauda sperm samples (from the same animal) are injected into control oocytes, and embryos are cultured to the indicated stages and subject to single-embryo RNA-Seq. Numbers associated with each stage give the number of embryos in the initial baseline dataset (excluding later RNA injection datasets – see **Table S8** for a complete listing of embryo numbers for the entire study). C) ICSI 2-cell stage dataset faithfully recapitulates the major wave of zygotic genome activation (ZGA). Scatterplot of average mRNA abundance (average tpm +1) for 18 hour embryos (pre-ZGA, x axis) vs. 28 hour embryos (post-ZGA, y axis), with significantly (adjusted p < 0.05) differentially-expressed genes shown as red and green dots. See also **Supplemental Figure S2**. D) Heatmaps of significant ZGA genes shown separately for caput- and cauda-derived embryos, as indicated. E–F) Scatterplots of average mRNA abundance (as in panel C) for Caput vs. Cauda embryos at 18 hours (E) and 28 hours (F), showing no significantly differentially-expressed genes.

Here, we use the dramatic remodeling of the sperm small RNA payload that occurs during epididymal transit to probe the function of sperm-delivered RNAs in the zygote. Sperm were obtained from the caput and cauda epididymis of the same animal, and matched zygotes were generated via intracytoplasmic sperm injection (ICSI). These zygotes were then developed to varying stages of preimplantation development for molecular characterization by single-embryo mRNA-sequencing, or transferred to surrogate mothers for further development. We find that caput-derived embryos significantly overexpress ~50 genes primarily encoding regulatory factors (RNA binding proteins and chromatin-associated factors), beginning at the 4-cell stage and persisting to the blastocyst stage of development. These preimplantation regulatory aberrations were accompanied by pleiotropic defects in implantation and post-implantation development, as caput-derived embryos transferred into surrogate mothers did not develop to term. Remarkably, defects in both early embryonic gene regulation, and in post-implantation development, could be rescued by microinjection of purified small RNAs from cauda epididymosomes into caput-derived ICSI zygotes. Finally, we show that microRNAs (or similarly-sized small RNAs), rather than the more abundant class of longer tRNA fragments, are the essential RNAs gained during epididymal transit. Together, our data reveal an essential role for epididymal RNA trafficking during post-testicular maturation of sperm, and identify a specific molecular program that is controlled by sperm-delivered microRNAs in preimplantation development.

## RESULTS

We take advantage of the dramatic remodeling of the sperm RNA repertoire that occurs during epididymal transit to interrogate the function of distinct RNA payloads in the zygote. Sperm were obtained from the caput and cauda epididymis of the same animal, and zygotes were generated via intracytoplasmic sperm injection (ICSI) using these paired samples (**Figure 1B**). For convenience, we refer throughout to embryos generated using cauda epididymal sperm as either “cauda-derived” ICSI embryos, or as “Cauda” embryos, to distinguish between caput/cauda as descriptors of location and Caput/Cauda as compact descriptors of the embryo generation protocol.

Focusing first on preimplantation development, we cultured Caput and Cauda embryos to varying developmental stages (2-cell, 4-cell, 8-cell, morula, and blastocyst) for molecular characterization by single-embryo RNA-Seq (Ramskold et al. 2012; Sharma et al. 2016). Importantly, both fertilization rates and blastocyst progression rates were identical for caput and cauda sperm (**Table S1**), ruling out the possibility that sperm obtained from the caput epididymis are grossly defective (aneuploid, etc.) sperm in the process of being resorbed.

**Table S1:** Fertilization and blastocyst progression rates for caput and cauda sperm.

|  |  |  |
| --- | --- | --- |
|  | Fertilization (progression of ICSI zygotes to 2-cell stage) | Blastocyst progression (% of 2-cell stage embryos progressing to blastocysts) |
| Caput | 77.8 +/- 1.9%<br>n=1155 | 60.3 +/- 4.7%<br>n=348 |
| Cauda | 75.6 +/- 2.7%<br>n=1181 | 62.2 +/- 4.9%<br>n=319 |

In addition to allowing us to generate embryos using caput sperm (which do not exhibit forward motility and are therefore not competent for in vitro fertilization), ICSI provides unprecedented temporal resolution for genome-wide analysis of early development, as the precise moment of fertilization is known (unlike in typical in vitro fertilization protocols). We therefore analyzed 2-cell stage embryos at two time points – 18 hours and 28 hours post–fertilization – which span the maternal-to-zygotic transition (**Table S2**). This dataset faithfully documents the process of the major wave of zygotic genome activation (ZGA), revealing 5654 genes that significantly change in abundance from 18 to 28 hours (**Figure 1C–D**). Additional embryos cultured for intermediate times provided a finer temporal view of this process (**Supplemental Figure S2**). Confident in our experimental pipeline, we next compared data for Caput and Cauda 2-cell stage embryos to identify any gene expression phenotypes induced by the developmental differences in the sperm small RNA epigenome. However, the source of sperm used for ICSI had no significant effect on mRNA abundance in either 18 or 28 hour 2-cell stage embryos (**Figure 1E–F**). While it is possible that subtle yet meaningful changes in mRNA abundance occur that are not captured by single-embryo RNA-Seq, we conclude that any such changes must be quantitatively modest at this stage of development.

### Overexpression of multiple epigenetic regulators in Caput embryos

We next turned to the single embryo RNA-Seq dataset for later stages of preimplantation development – 4-cell, 8-cell, morula, and blastocyst (**Tables S3–4**) – focusing first on the 4-cell stage. Here, we find a strong signature of differential gene expression, with 37 genes exhibiting significantly increased mRNA abundance in Caput embryos, relative to matched cauda-derived embryos (**Figures 2A–B**). These genes primarily encode regulatory factors, including RNA binding proteins (Hnrnpab, Hnrnpu, Pcbp1, Eif3b, Yrdc, Ythdf1, Srsf2, and Ybx1) and chromatin-associated factors (Smarcd2, Smarca5, Smarcc1, Trim28, and Ezh2). Not only were these genes upregulated at the 4-cell stage, but they (along with other RNA and chromatin regulatory genes – Grsf1, Alyref, etc. – that changed, but were not statistically significant, in the 4cell dataset) remained upregulated in 8-cell, morula, and blastocyst-stage embryos (**Figures 2C–D, Supplemental Figures S3–4**). The consistent behavior of these genes throughout preimplantation development provides extensive biological replication of this gene signature, giving us a high–confidence set of target genes affected by some aspect of post–testicular sperm maturation. In addition to the common gene expression signature observed across preimplantation development, a small number of genes were preferentially affected specifically at the blastocyst stage, with decreased levels of *Upp1, Utf1*, and *Tdgf1*, and increased levels of *Gata6*, in Caput blastocysts (**Figure 2D, Supplemental Figure S4B**). Together, our preimplantation RNA-Seq data reveal robust effects of sperm maturation on expression of a group of RNA and chromatin regulators, with these early expression changes potentially contributing to later effects on signaling and gene regulation in the blastocyst.

**Figure 2.**
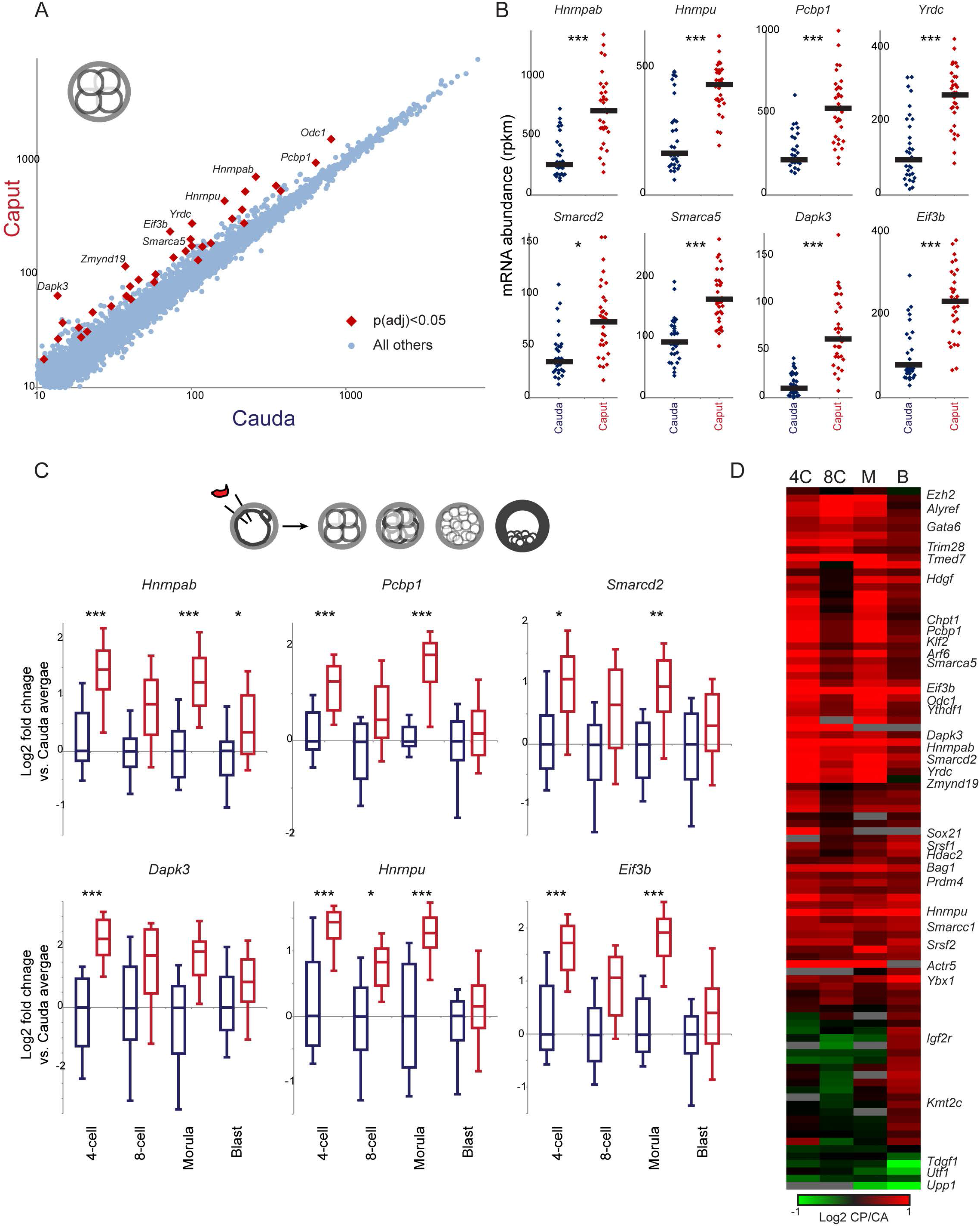
Effects of post-testicular sperm maturation on preimplantation gene regulation. A) Scatterplot of 4-cell stage gene expression for Cauda vs. Caput embryos. Data are shown for all genes with a median expression level of at least 10 tpm across the entire dataset, and significant differences (here, showing 37 genes with t-test p value < 0.05, adjusted for multiple hypotheses) in gene expression are indicated. B) Data for every individual 4-cell embryo for eight genes overexpressed in Caput embryos. Black bars show median expression for each group of embryos. *, **, and *** show comparisons with adjusted p values below 0.1, 0.01, and 0.001, respectively. C) Gene expression differences at the 4-cell stage persist into later developmental stages. Here, box plots (box: median and 25^th^/75^th^ percentiles, whiskers: 10^th^ and 90^th^ percentiles) show log_2_ of expression level normalized relative to the median value in matched cauda-derived embryos of the relevant developmental stage. D) Heatmap of all 95 genes exhibiting a difference in expression (p_adj_ < 0.1) at any stage of development. Data are expressed as log_2_ ratio (Caput/Cauda).

### Caput–derived embryos exhibit multiple defects in post-implantation development

Interestingly, the significant regulatory changes observed in early embryos do not impact overall preimplantation development – we achieved identical rates of fertilization and blastocyst progression for ICSI using caput and cauda sperm (**Table S1**). This raises the question of whether the significant molecular alterations observed in early development might result in any overt later phenotypes, such as defects in postimplantation development, or altered physiology in resulting offspring. To address this question, we generated Caput and Cauda embryos by ICSI using paired sperm samples as described above, cultured the embryos to the 2-cell stage, and then surgically transferred 10–20 embryos into pseudo-pregnant surrogate mothers (**Figure 3A**). Although we initially set out to characterize phenotypes in offspring of these experiments, to our surprise none of the females implanted with Caput embryos ever gave birth (**Figure 3B**). This was not a defect in our embryo transfer procedure, as we readily obtained healthy litters from matched transfers of cauda-derived ICSI embryos carried out under identical conditions. Given that Caput embryos are perfectly capable of progression to the blastocyst stage (**Table S1**), we conclude that caput sperm carry a functional haploid genome (e.g. they are not aneuploid, or heavily oxidatively damaged), and instead that caput sperm may lack some epigenetic factor required for normal postimplantation development.

**Figure 3.**
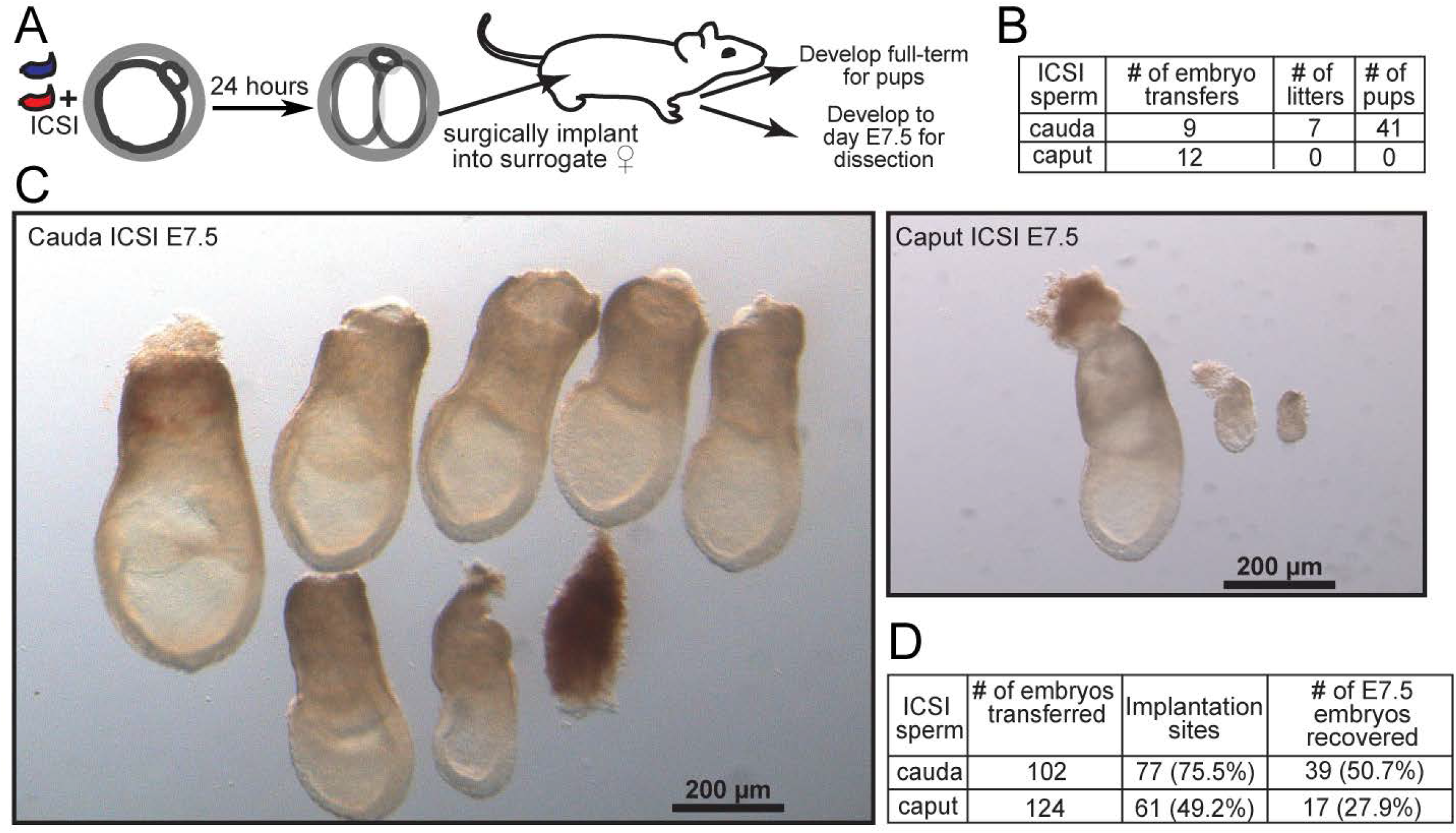
caput-derived embryos fail during early post-implantation development. A) Schematic representation of embryo transfer experiments. ICSI embryos were cultured to the 2-cell stage, then 15–20 embryos were transferred into an oviduct of a pseudo-pregnant Swiss Webster female. B) Success rates for Caput and Cauda embryos allowed to develop to term. Note that Caput embryos were also transferred to recipient females in additional experiments below, with a total success rate of 0/20 across all experiments in this study. C) Images of all embryos dissected from females transferred with Cauda or Caput embryos. A greater fraction of Cauda embryos implanted and produced normally-developing embryos. The left panel shows six normal, and one abnormal, Cauda embryos. In contrast, fewer implantation sites were observed for the Caput embryos, and the few implantation sites observed were either undergoing resorption or carried abnormally developing embryos. The right panel shows one normal embryo and two abnormal ones. D) Quantitation of all embryo transfer dissections. caput-derived embryos exhibit lower implantation rates, higher resorption rates, and (not quantified but shown in panel C) poor growth in the peri-implantation period. Note that percentages associated with number of E7.5 embryos recovered are relative to number of implantation sites, not total number of embryos transferred.

At what stage do Caput embryos fail during post-implantation development? We initially noted that recipient females carrying Caput embryos do not exhibit the increase in size typical of gravid females, suggesting that the defect occurs relatively early in pregnancy. We therefore repeated ICSI with caput and cauda sperm and transferred 2-cell stage embryos as before, but rather than allow the embryos to go to term we dissected the embryos from the uteri at embryonic day 7.5 (E7.5). **Figure 3C** shows all embryos recovered from a typical pair of embryo transfers, with Cauda embryos yielding a number of large, normally-developing embryos, along with a few defective embryos. In contrast, we obtained many fewer Caput embryos at this stage of development, and the majority of these exhibited clear growth defects. We quantified several features of peri-implantation development, finding that the decrease in the number of healthy caput-derived embryos was a result of defects at multiple stages: Caput embryos exhibited fewer implantation sites in the recipient uterus (for the same number of transferred embryos), a higher fraction of embryos were resorbed, and the small number of developing embryos more often exhibited gross delays in developmental progression (**Figure 3D**). Taken together, these results demonstrate that, in our hands, Caput embryos exhibit a pleiotropic defect during or soon after implantation (see Discussion).

### Molecular and developmental characterization of embryos generated using testicular spermatozoa

The failure of Caput ICSI embryos to develop to term is quite surprising in light of the fact that embryos generated using even more immature testicular sperm, or even round spermatids, are viable – this is a common method of assisted reproduction in humans (Greco et al. 2005; Tanaka et al. 2015). We readily confirmed this finding under our experimental conditions, successfully producing offspring from ICSI embryos using testicular spermatozoa (3/4 transfers yielding offspring), in contrast to 0/3 litters for a matched set of Caput embryo transfers. Given the ability of testicular sperm ICSI embryos to develop to term, we tested the prediction that the preimplantation regulatory aberrations observed in Caput embryos would not occur in Testicular embryos. Intriguingly, the altered gene expression program observed in Caput embryos was indeed absent in embryos generated using testicular spermatozoa (**Figure 5A, Supplemental Figure S5**), confirming that both testicular sperm and cauda epididymal sperm are capable of supporting similar embryonic development from fertilization to birth, while caput sperm – uniquely – cause the regulatory and developmental aberrations documented in **Figures 2–3**. These data suggest two likely hypotheses: 1) that sperm isolated from the caput epididymis represent a developmental dead end resulting from defects such as, eg, DNA damage, or 2) that epigenetic dynamics during early post-testicular development create a transiently-defective sperm population that is subsequently corrected during further epididymal transit.

### Small RNAs gained during epididymal transit rescue preimplantation defects in caput-derived embryos

As Caput and Cauda ICSI embryos are identically successful through blastocyst development (**Table S1**), we set out to test the hypothesis that epigenetic differences between caput and cauda sperm are responsible for the preimplantation molecular alterations, and the defective post-implantation development, of caput-derived embryos.

Cytosine methylation patterns are largely unchanged following their establishment during primordial germ cell development (Feng et al. 2010) – we have confirmed in whole genome surveys that genes in the Caput gene expression program are not differentially-methylated between caput and cauda sperm (unpublished data) – and the histone-to-protamine transition is already essentially complete in testicular spermatozoa (Gaucher et al. 2010). We therefore focused on the possibility that the dramatic remodeling of the sperm RNA payload that occurs during epididymal maturation (**Figure 1A**) could be responsible for the effects we see here. To test this, we obtained cauda-specific small RNAs by purifying 18–40 nucleotide RNAs from extracellular vesicles – epididymosomes – isolated from the cauda epididymis (Sharma et al. 2016). We then generated three sets of embryos: Cauda ICSI embryos, control Caput ICSI embryos injected with H3.3-GFP mRNA (to visualize successfully-injected embryos), and Caput ICSI embryos injected with cauda–specific small RNAs (as well as H3.3-GFP) three hours after fertilization (**Figure 5A**). Resulting embryos were cultured to the blastocyst stage and characterized by RNA-Seq (**Table S5**).

Consistent with the first two replicate datasets and with data from earlier developmental stages (4-cell, etc.), this third blastocyst dataset again reveals upregulation of the same group of regulatory genes – *Hnrnpab, Eif3b, Pcbp1, Gata6*, etc. – in caput-derived embryos compared to Cauda embryos (**Figure 5B**). Strikingly, injection of cauda-specific small RNAs was sufficient to rescue the gene expression defects of Caput embryos, offsetting the overexpression of these regulatory genes (**Figure 5B**). More globally, cauda-specific small RNAs were able to almost completely restore Cauda-like gene expression in Caput blastocysts (**Figure 5C**). The complete suppression of the Caput preimplantation gene expression program by small RNA injections essentially rules out the possibilities that this altered regulatory program results from 1) defects in caput sperm such as aneuploidy or oxidative DNA damage, 2) cytosine methylation or chromatin differences between caput and cauda sperm, or 3) the absence of various proteins normally delivered to sperm during epididymal maturation (Sullivan et al. 2007; Krapf et al. 2012).

What is the identity of the small RNAs responsible for rescuing the preimplantation regulatory defects in Caput embryos? As detailed extensively elsewhere ((Sharma et al. 2016) and Sharma *et al*, accompanying manuscript, summarized in **Figure 1A and Supplemental Figure 1**), caput sperm gain multiple small RNAs as they continue their transit through the epididymis. cauda-specific RNAs most notably include several abundant tRFs (tRF-Val-CAC, tRF-Val-AAC), as well as ~100 microRNAs, most of which are encoded in multi-microRNA genomic clusters (as in, eg, the X-linked miR-880 cluster, or the “oncomiR”17–92 cluster). To broadly separate the effects of cauda-specific tRFs from cauda-specific microRNAs, we gel-purified 18–26 nt and 26–40 nt cauda epididymosomal RNAs to enrich for microRNAs and tRFs, respectively (**Figures 6A–B**). It is important to note that although this procedure greatly enriches the indicated RNA species (free of associated binding proteins), these preparations nonetheless represent complex RNA populations, as for example the “microRNA” fraction includes low levels of a variety of relatively short (<26 nt) tRNA fragments (see Discussion). Purified small RNA populations were microinjected into caput-derived ICSI embryos as described above, and embryos were cultured to either the 4-cell stage or to the blastocyst stage for single-embryo RNA-Seq (**Tables S6–7**).

These data once again confirmed the robust overexpression of *Hnrnpab* and various other genes involved in RNA and chromatin metabolism in Caput embryos. This Caput-specific gene expression program was unperturbed by microinjection of cauda-specific tRFs, both for 4-cell stage embryos and for blastocysts (**Figure 6C** and **Supplemental Figure S6B**). Instead, we find that the suppression of the Caput preimplantation gene expression program by cauda-specific small RNAs could be recapitulated via microinjection of cauda-specific microRNAs (**Figures 6C–D and Supplemental Figure S6**). This was readily apparent for any individual Caput-specific target gene (**Figure 6C, Supplemental Figure S6B**), and more generally we find that a Cauda-like gene expression program was globally restored in caput sperm-derived embryos following microRNA injection (**Figure 6D, Supplemental Figure S6C**). Repression of Caput–induced genes was specific to cauda-specific microRNAs, as cauda-specific tRFs had no effect on these genes (**Figures 6E–F**). Taken together, these data demonstrate that the microRNAs gained by caput sperm as they further transit the epididymis are required to support normal preimplantation gene regulation.

### cauda-specific small RNAs rescue development of caput-derived embryos

Finally, as cauda-specific small RNAs were able to rescue the preimplantation molecular defects in caput-derived embryos, we sought to determine whether these RNAs could also rescue the post-implantation defects of these embryos. We therefore generated the same three sets of embryos described in **Figure 5A** – Cauda, Caput, and Caput+RNA – which were then transferred into recipient females as in **Figure 3**. As observed in **Figure 3**, we did not obtain any live births from the Caput embryos – across three different experiments we obtained 0 litters from 20 Caput embryo transfers (in contrast to 7/9 for Cauda embryos, or 3/4 for Testicular embryos). Astonishingly, microinjection of cauda-specific small RNAs suppressed the developmental lethality observed in the Caput embryos, as we obtained litters for 4 out of 7 embryo transfers (**Figure 7**). We conclude that the small RNAs that are gained as sperm transit from the caput to the cauda epididymis are essential to support proper development. Moreover, the ability of small RNAs to rescue the development of embryos fertilized using caput epididymal sperm provides further evidence that these sperm are not genetically defective, but are simply epigenetically immature.

## DISCUSSION

Taken together, our data demonstrate that the dynamic remodeling of the small RNA payload of sperm that occurs during post-testicular maturation has dramatic functional consequences in the early embryo, and define a novel function for the epididymal maturation process beyond the established acquisition of sperm motility and competence for fertilization. We show that paternally-transmitted small RNAs function during preimplantation development to downregulate a set of genes encoding RNA-binding proteins and chromatin-associated factors, and that proper acquisition of small RNAs by maturing sperm is essential for offspring development. Importantly, several lines of evidence rule out the hypothesis that sperm obtained from the caput epididymis represent defective sperm in the process of apoptosis or resorption, as 1) fertilization rates and blastocyst progression are identical for Caput and Cauda embryos, and 2) the developmental defects of Caput embryos can be rescued simply by small RNA microinjection.

We are aware that our findings appear to contradict previous reports documenting successful generation of full-term offspring from Caput ICSI embryos in mouse (Suganuma et al. 2005) and (with extremely low ICSI success rates) in rat (Said et al. 2003). Focusing on the more comparable mouse experiments, we consider several potential explanations for this discrepancy. First, our studies differ in strain background – we used FVB rather than 129Sv animals – and it is plausible that this affects the ability of Caput embryos to implant. Second, Suganuma *et al* obtained sperm from 6-10 month old males, whereas we use 10-12 week old males. Third, the precise anatomical dissection of the caput epididymis could differ between our preparations – given that both testicular sperm and cauda sperm support full development in our hands as well as countless other reports, a caput epididymis dissection that included earlier or later sperm populations could result in greater developmental success of “caput sperm” ICSI embryos. That said, the comparable success rate for Caput and Cauda embryos reported by Suganuma *et al* makes this idea unlikely. Fourth, our protocols differ significantly in the embryos transferred to females, as Suganuma *et al* transferred morula/blastocyst stage embryos to the ampulla, whereas we transfer 2-cell stage embryos. It is plausible that multiple days of diapause experienced by blastocyst-stage Caput embryos as they transit the oviduct could potentially help overcome the deficits we observe in Caput embryos. Finally, we note that ICSI is typically performed with sperm heads, but sperm head isolation protocols differ in detail – Suganuma *et al* generated sperm heads by drawing sperm into the injection pipette, applying a few piezo pulses, and recovering the sperm head for injection. In rats, Said *et al* generated sperm heads by sonicating intact sperm. In contrast, we generate sperm heads en masse by drawing sperm repeatedly through a fine-gauge needle (Methods), and sperm heads from the resulting suspension are then picked for ICSI. It is likely that these protocols differ in the extent to which the injected sperm heads retain associated sperm components such as the midpiece, and it is therefore almost certain that the RNA contents differ between caput sperm head preparations.

Whatever the reason for this discrepancy, we demonstrate here that injection of cauda epididysomal small RNAs reproducibly and completely rescues both the defective development and molecular gene expression phenotype exhibited by embryos fertilized with our caput sperm head preps. This is clearly most consistent with a model in which sperm small RNAs gained during epididymal transit have important regulatory functions during early embryonic development.

### A functional role for microRNA remodeling in the epididymis

Sperm undergo at least two dramatic alterations to their small RNA payload during the process of post-testicular maturation in the epididymis – a global switch from a piRNA-dominated payload in testicular sperm to a tRF-dominated payload in the epididymis, and a transient destruction or loss of a subset of microRNAs in caput sperm that is then countered by re-gain of these microRNAs in the cauda epididymis (**Supplemental Figure S1**). Two lines of evidence demonstrate that it is this latter process – the transient loss of genomically-clustered microRNAs – that causes the developmental aberrations in embryos generated using caput epididymal sperm. First, the robust overexpression of various regulatory genes is observed specifically in embryos generated using caput epididymal sperm, whereas embryos generated using testicular spermatozoa and cauda epididymal sperm exhibit much more similar gene expression programs (**Figure 4**). Moreover, in contrast to Caput embryos, we readily obtained offspring from both Cauda and Testicular ICSI embryos. As the loss of clustered microRNAs such as the miR-880 cluster is a transient feature specific to caput sperm – these microRNAs are abundant in all testicular sperm populations analyzed from primary spermatocytes to mature spermatozoa, and are similarly-abundant in cauda sperm – the strong similarity between Testicular and Cauda embryos pointed to the possibility that these microRNAs might play a role in early embryonic gene regulation. Second, fractionation of the cauda-specific small RNA payload based on RNA length confirmed that microRNAs (or, less likely, relatively low-abundance fragments of mature tRNAs that are shorter than the primary population of 28-36 nt tRNA cleavage products) are responsible for supporting normal gene expression in Cauda embryos, as reintroduction of this purified RNA population was able to almost completely suppress the overexpression of regulatory genes in caput-derived embryos (**Figure 6**). Moreover, in ongoing studies we have been able to obtain viable offspring from Caput ICSI embryos injected with this microRNA fraction (2/4 transfers). We note that although our gel-purified “microRNA” fraction includes additional RNA species including relatively short tRNA fragments, the absence of the Caput regulatory aberrations in embryos generated using testicular sperm – which have yet to gain appreciable tRFs, but which express the microRNAs in question – strongly argues that it is indeed the microRNAs in this fraction that rescue the developmental deficits in Caput embryos.

**Figure 4.**
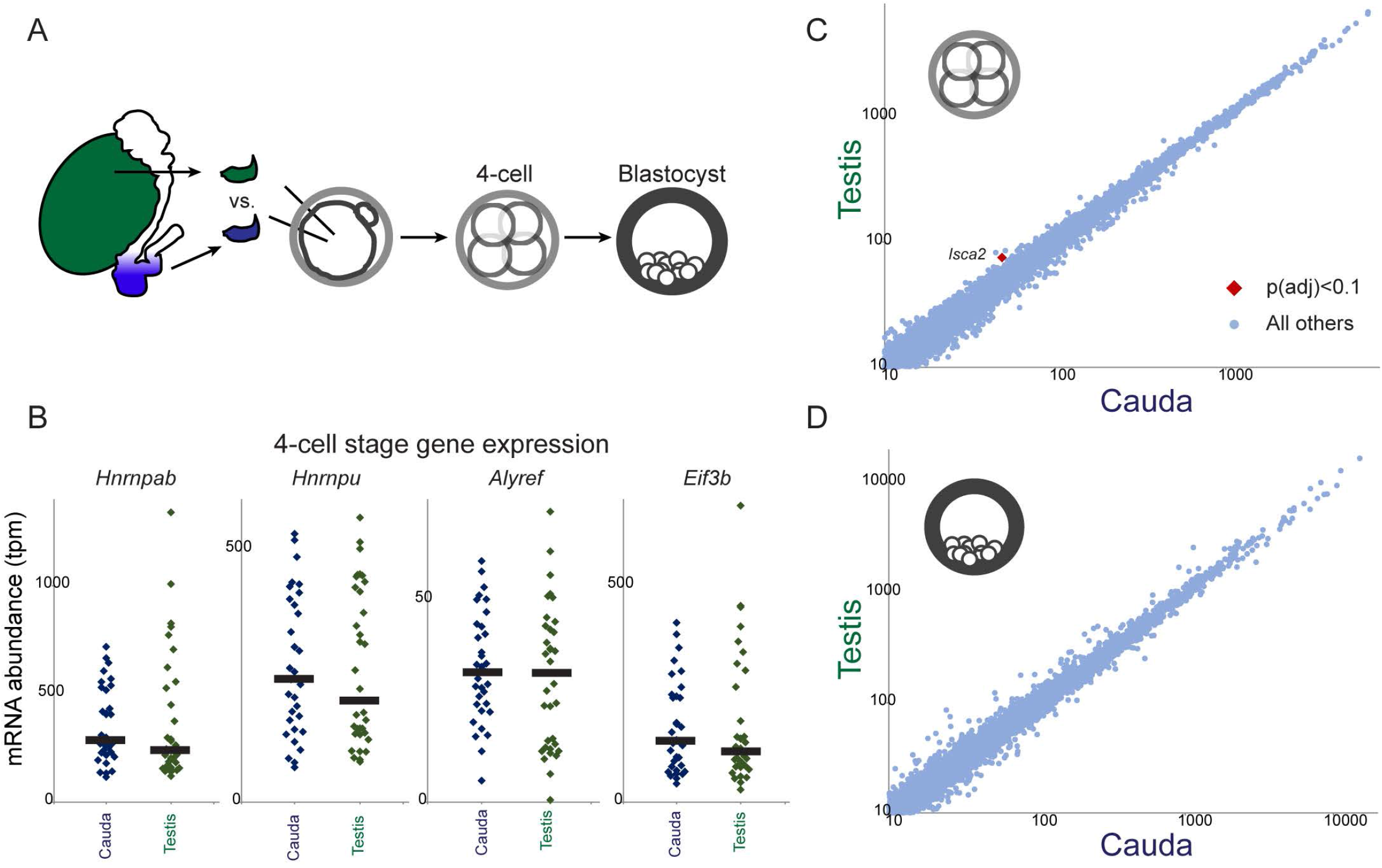
Molecular analysis of testicular sperm-derived preimplantation embryos. A) Schematic of testicular sperm ICSI experiments. B) Dot plots showing examples of key Caput-regulated genes in individual 4-cell stage Cauda (n=33) and Testicular (n=34) embryos. Black bars show median expression level for each condition. C-D) Scatterplots comparing mRNA abundance (all genes with median tpm > 10) between Cauda and Testicular embryos at the 4-cell (C) and blastocyst (D) stages. See also **Supplemental Figure S5** for morula-stage data.

**Figure 5.**
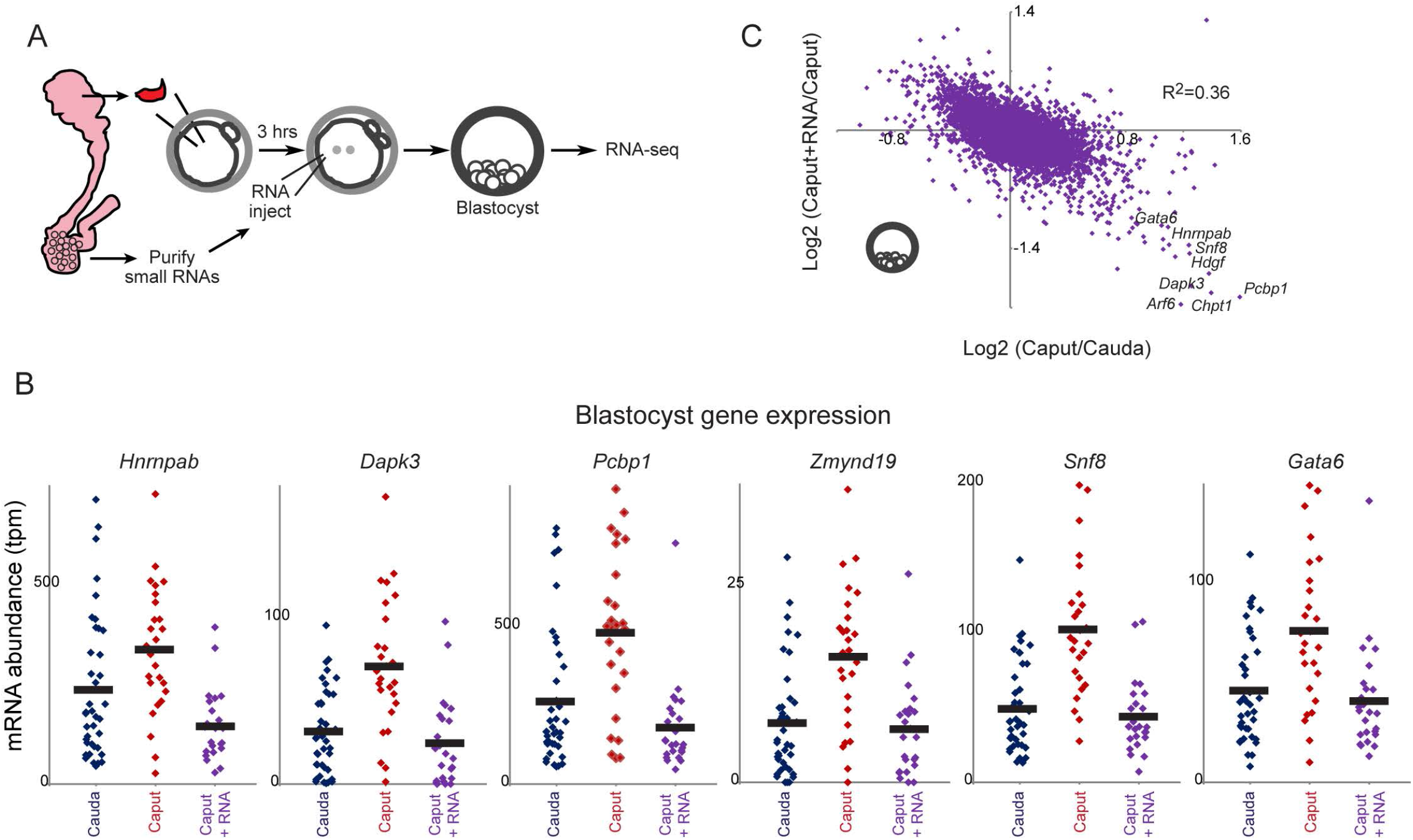
cauda-specific small RNAs rescue development of caput-derived embryos. A) Schematic representation of cauda epididymosomal RNA purification and microinjection. ICSI zygotes were generated by injection of caput or cauda sperm. Three hours later, Caput embryos were microinjected either with gel-purified small (1840 nt) RNAs obtained from cauda epididymosomes, along with H3.3-GFP mRNA as a tracer for successful injections, or were injected with H3.3-GFP alone. Resulting zygotes were cultured to the blastocyst stage for RNA-Seq. B) cauda-specific RNAs repress genes overexpressed in caput-derived embryos. Dots show individual embryo RNA-Seq data for Cauda (n=39), Caput (n=26), or Caput+RNA (n=25) blastocysts. In every case, injection of cauda-specific small RNAs returns expression of these genes to the Cauda baseline. C) Scatterplot of Caput effects on mRNA abundance (x axis, log2 fold change Caput/Cauda) vs. effects of small RNA injection (y axis, log2 fold change Caput+RNA/Caput). Dots in lower right corner are overexpressed in Caput embryos, and downregulated upon microinjection of cauda-specific small RNAs.

**Figure 6.**
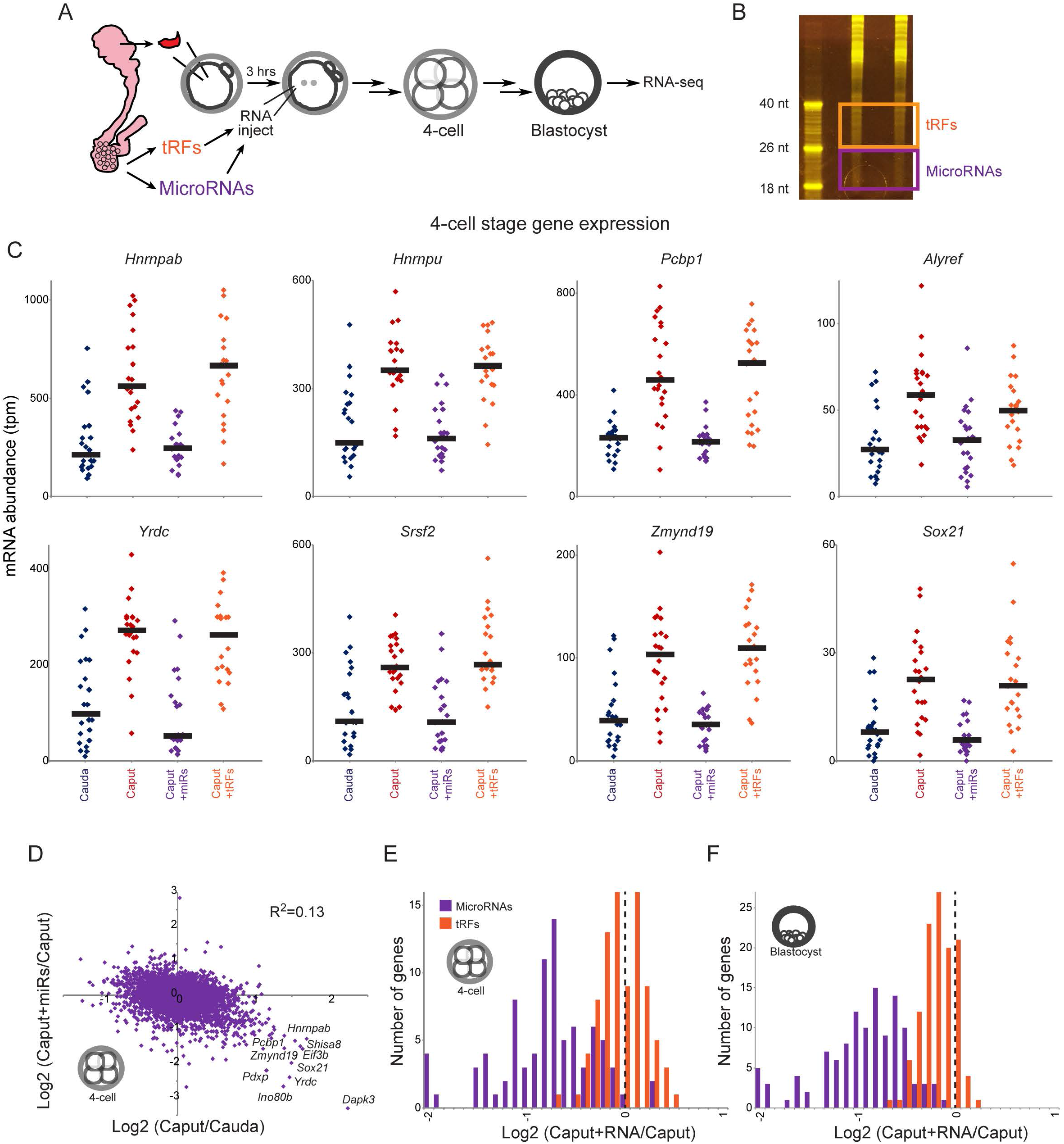
cauda-specific microRNAs, but not tRFs, rescue Caput preimplantation defects. A) Schematic representation of microinjection experiment. As in **Figure 5A**, but two separate small RNA populations were gel-purified – 18–26 nt RNAs are enriched for microRNAs (but include additional molecular species including short 5’ and 3’ tRFs), while 26–40 nt RNAs are primarily comprised of tRNA fragments. B) Purification of epididymosomal RNAs. Acrylamide electrophoresis of two replicate total RNA samples purified from cauda epididymosomes. Boxes show the boundaries used for gel-purification of microRNAs and tRFs. C) mRNA abundance for individual 4-cell stage Cauda (n=23), Caput (n=23), Caput+microRNA (n=22), and Caput+tRF (n=21) embryos, for representative Caput-upregulated genes. In all cases, cauda-specific “microRNAs” returned mRNA abundance to Cauda levels, while tRFs had little to no effect on these target genes. See **Supplemental Figure S6B** for blastocyst-stage data. The clear difference between these matched RNA injections – both RNA samples should be free of contaminating proteins, and should include similar levels of any potential contaminants such as leftover acrylamide – demonstrates that the rescue activity is specific to 18–26 nt cauda epididymosomal small RNAs. D) Scatterplot comparing Caput effects (x axis) and microRNA effects (y axis) on mRNA abundance in 4-cell stage embryos. See **Supplemental Figure S6C** for blastocyst stage dataset. E–F) Histograms of microRNA and tRF effects on Caput-upregulated genes. In both cases, data are shown for mRNAs which exhibit a log_2_ fold change of at least 0.5 in Caput vs. Cauda embryos (p<0.01, uncorrected for multiple hypothesis testing to include a greater number of genes – results are even more dramatic for the smaller subset of genes significant after multiple hypothesis-correction). For both 4-cell stage (E) and blastocyst-stage (F) data, cauda-specific microRNAs cause downregulation of Caput-induced genes, while tRFs have little to no effect on these genes (KS p value comparing microRNA and tRF effects < 10^−15^ for both datasets).

**Figure 7.**
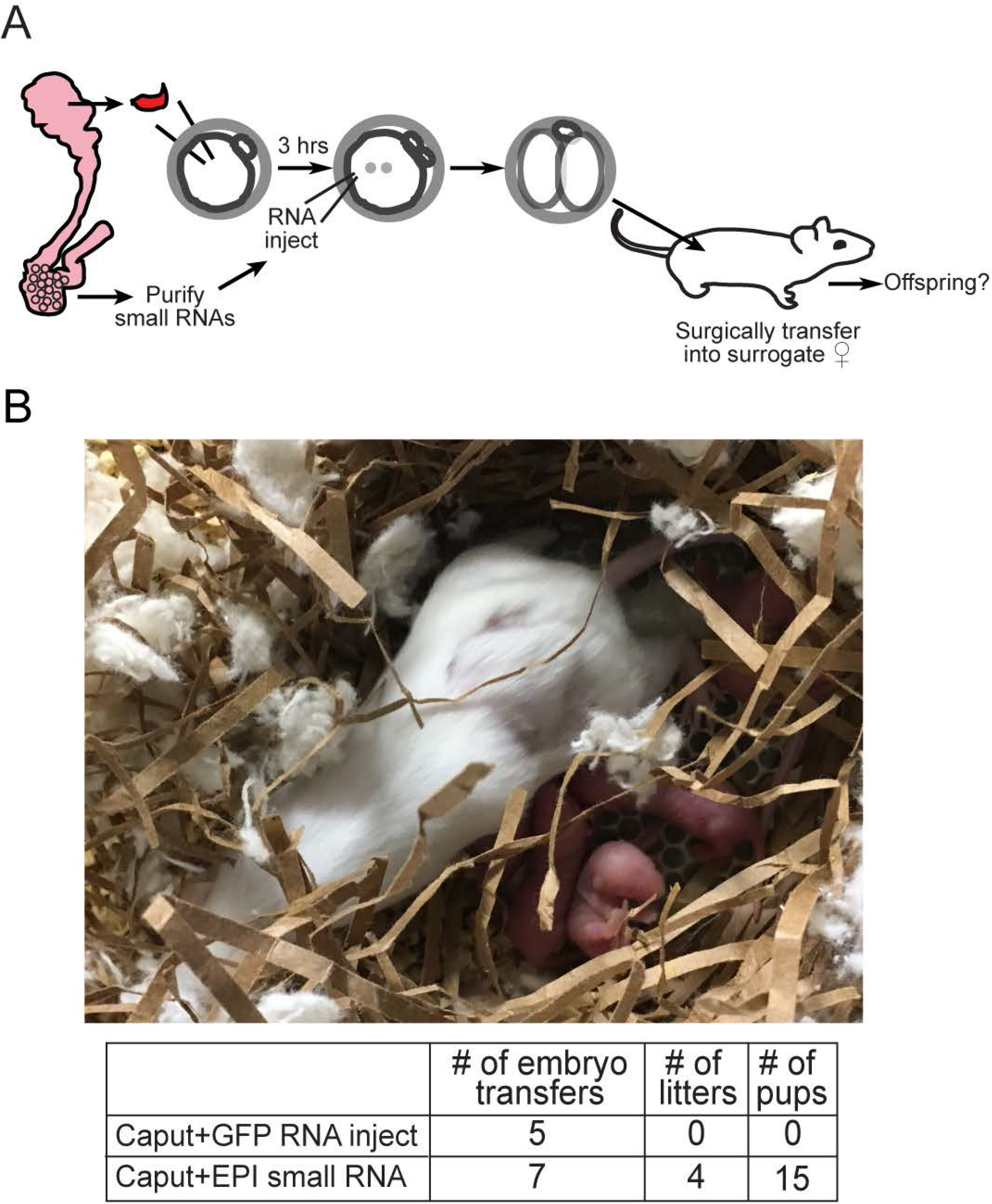
cauda-specific small RNAs rescue development of Caput embryos. A) Experimental schematic. Small RNA injections were carried out as in **Figure 5A**, but ICSI embryos were transferred at the 2-cell stage into the oviduct of pseudo-pregnant females as in **Figure 3**. B) Image shows example of a successful litter generated from caput-derived embryos microinjected with cauda RNAs. Table underneath lists success rates, as in **Figure 3B**.

These data identify a functional consequence for a surprising epigenetic remodeling event – transient elimination of a group of genomically-clustered microRNAs – recently documented to occur during post-testicular sperm maturation (Sharma et al. 2016). This process bears a curious similarity to the global erasure and reestablishment of cytosine methylation patterns that occurs following fertilization, and again during primordial germ cell development, in mammals (Feng et al. 2010). The purpose of this microRNA turnover is unclear, although as caput sperm are not transferred to the female during natural mating and in any case are incapable of fertilization – caput sperm do not exhibit forward motility – we do not anticipate that this process normally plays any role in control of offspring phenotype. Nonetheless, together with a prior report demonstrating that defective blastocyst progression of embryos generated from *Dicer* cKO germ cells can be rescued with sperm RNAs (Yuan et al. 2016), our findings provide very strong additional evidence that the mammalian sperm RNA payload is functional upon introduction to the zygote.

The identity of the small RNA(s) responsible for repression of the various RNA and chromatin regulatory genes identified here remains to be determined. Roughly ~100 microRNAs present at >50 transcript per million abundance in cauda sperm are at least 2–fold less abundant in caput sperm (Sharma et al. 2016), and extensive examination of predicted microRNA targets reveals no strong single candidate microRNA as a common regulator of the various Caput-upregulated target genes. We note two broad possible explanations for the observed Caput gene expression profile. First, given the number of differentially-abundant microRNAs, it is plausible that multiple distinct microRNAs each repress expression of a small number of relevant target genes – that, for example, miR–17 represses *Grsfl*, miR-881 represses *Hnrnpu*, etc. Alternatively, one specific microRNA-mediated regulatory event could cause a broad downstream regulatory program, as for example repression of a single signaling protein or a transcription factor could result in altered expression of the ~50 target genes observed here. Beyond microRNA targeting of some upstream regulator, the altered gene expression program could be a physiological response to a global cellular defect such as, for example, decreased nuclear pore formation, altered nuclear exosome activity, etc. Given that the Caput-specific gene expression program is comprised of a strikingly coherent set of chromatin and RNA regulators, we favor the latter possibility – that the Caput gene expression program represents a cellular response to some upstream cellular defect – but this remains to be identified.

Together, our data demonstrate that paternally-transmitted small RNAs are functional in the early embryo and are essential for proper development. It will be interesting to determine the direct targets of sperm microRNAs in the zygote, the course of events that leads to downregulation of various regulatory factors during early development, and the specific molecular events responsible for facilitating proper postimplantation development. Finally, although embryos generated using immature testicular sperm are viable in mouse and humans, given that testicular sperm carry a very distinct small RNA repertoire from that of cauda sperm we suggest that the phenotypic consequences of ICSI bear further scrutiny, and our studies provide a possible interventional framework for offsetting potential issues.

## ACKNOWLEDGEMENTS

We thank members of the Rando lab for critical reading of the manuscript. Research was supported by NIH grants R01HD080224 and DP1ES025458 (OJR) and R01HD083311 (JAR–P). CCC was supported by a Helen Hay Whitney Foundation Postdoctoral Fellowship.

## MATERIALS AND METHODS

### Intracytoplasmic Sperm Injection (ICSI)

The caput and cauda epididymis of 8–12 week old FVB/NJ mice were dissected separately into PBS. Sperm were then released by making an incision in the epididymal tissue followed by squeezing to release the contents. To detach sperm heads from the tails (both caput and cauda), after collecting the sperm in PBS in an eppendorf tube, the sperm were spun at 10,000 × g and washed once with PBS. The sperm were then resuspended in 500 μl PBS and then drawn through a 26G needle on a 1ml syringe between 20–30 times. The shearing force from being drawn in and out of the needle removes the sperm head from tail for the majority of sperm. The sperm were then washed twice in Nuclear Isolation Media (NIM) 1% PVA and finally resuspended in 100–500 μl NIM 1% PVA for use in ICSI.

Females were superovulated by an intraperitoneal (i.p) injection of pregnant mare’s serum gonadotropin (PMSG; 5 IU) followed by an i.p. injection of human chorionic gonadotropin (hCG; 5 IU) 48 hours later. Eggs were then collected from the oviducts of the females 13–18 hours later by placing the dissected ampulla of the oviduct into KSOM containing 3 mg/ml hyaluranidase to digest the cumulus cells away from the eggs. After several minutes in hyaluranidase the eggs were washed 4–5 times in KSOM, finally being placed in KSOM in a 37°C incubator until injected.

For ICSI, plates were made with drops of NIM with 1% Polyvinyl alcohol for washing the injection needle, drops of NIM 1% PVA with sperm, drops of FHM with 0.1% PVA for the eggs to be added to for injection, and finally covered with mineral oli. 15–25 eggs at a time were placed into the FHM 0.1% FHM drops of the injection plate for subsequent injection. Sperm heads were then picked and injected into the eggs. After completion of the ~20 injections, the injected eggs were maintained at room temperature for 5 minutes, washed 4 times in KSOM, and then cultured in KSOM in a 37°C incubator in 5% CO_2_, 5% O_2_, for development. The process was repeated for a total of 100–200 injections per day.

18, 24, and 28 hours post-injection, 2-cell embryos were collected into TCL buffer with 1% β–mercaptoethanol and then stored at –80°C for processing into single embryo RNA-Sequencing libraries. 4-cell, 8-cell, and morula embryos were collected at 46, 60 and 75 hours respectively. Blastocysts were collected 4–5 days after ICSI determined by the morphology of a typical mid-blastocyst stage.

### Epididymosome Small RNA Isolation and Injection

Epididymosomes were prepared and RNA was extracted as previously described (Belleannee et al. 2013; Sharma et al. 2016). In brief, luminal contents of the cauda epididymis were centrifuged at 2000 × g to pellet sperm, and the resulting supernatants were centrifuged at 10.0 × g for 30 minutes to get rid of cellular debris. Next the supernatants were subjected to an ultracentrifugation at 120,000 × g at 4 °C for 2 h (TLA100.4 rotor; Beckman). Pellets were washed in cold PBS and subjected to a second ultracentrifugation at 120,000 × g at 4 °C for 2 hours. These pellets were then resuspended in 50 μl of PBS and used for RNA extraction.

For epididymosome RNA extraction, the total volume of the samples were adjusted to 60 μl with water. 33.3 μl of lysis buffer (6.4 M Guanidine HCl, 5% Tween 20, 5% Triton, 120 mM EDTA, and 120 mM Tris pH 8.0), 3.3 μl ProteinaseK (>600 mAU/ml, Qiagen 19131), and 3.3 μl water were added to the samples. The samples were then incubated, with shaking, at 60 °C for 15 minutes on an Eppendorf thermomixer.

One volume of water (100 μl) was then added and the sample transferred to a phase lock column (5 PRIME). For phase separation 200 μl of TRI Reagent (MRC inc) and 40 μl BCP (1-bromo-2 chloropropane, MRC inc) were added. The samples were then mixed by inversion 10–15 times, followed by centrifugation at 14,000 RPM for 4 minutes. A second addition of TRI reagent and BCP was performed to further purify the RNA. The aqueous phase was then removed and transferred to a low binding RNase/DNase free microcentrifuge tube (MSP); followed by the addition of 20 μg of glycoblue (Ambion) and 1 volume (~200 μl) of Isopropanol. The RNA was then precipitated for 30 minutes or greater at –20°C, followed by centrifugation at 14.0 RPM for 15 minutes at 4°C, and one wash with 70% cold Ethanol followed by centriguation at 14,000 RPM for 5 minutes at 4°C. Finally, the RNA was reconstituted in 10 μl water for gel size selection.

Total RNA was combined with an equal volume of Gel Loading Buffer II (Ambion), loaded onto a 15% Polyacrylamide with 7M Urea and 1X TBE gel, and run at 15W in 1X TBE until the dye front was at the very bottom of the gel (~25 minutes for Criterion minigels). After staining with SYBR Gold (Life Technologies) for 7 minutes, and destaining in 1X TBE for 7 minutes, gel slices corresponding to 18–40 nucleotides were then cut from the gel. Gel slices were then ground (using a pipette tip or plastic pestle) and 750 μl of 0.3 M NaCl–TE pH 7.5 was added and incubated with shaking on a thermomixer overnight at room temperature. The samples were then filtered using a 0.4 μm Cellulose Acetate filter (Costar) to remove gel debris. The eluate was transferred to a new low binding microcentrifuge tube and 20 μg of glycoblue and 1 volume of Isopropanol (~700 μl) were added. Finally, samples were precipitated for 30 or more minutes at –20 °C.

The resulting small RNA preps are almost certainly free of ribonucleoprotein complexes, as they are purified from acrylamide gels. Although they may contain other impurities (traces of acrylamide, etc.), the clear distinction between the “microRNA” and “tRF” preps in microinjection studies (**Figure 6, Supplemental Figure S6, Supplemental Tables S6–7**) strongly argues that the relevant activity of these preps is in the specific RNA populations purified.

Three hours following ICSI fertilization, zygotes were transferred to an injection plate with FHM medium containing 0.1% PVA, and subjected to micromanipulation. Caput zygotes were microinjected with either H3.3-GFP mRNA (caput control) or H3.3-GFP mRNA plus 1 ng/μl epididymosomal gel-purified small RNAs (18–40 nts). RNA injections were carried out using a Femtojet (Eppendorf) microinjector at 100 hPa pressure for 0.2 seconds, with 7 hPa compensation pressure. After the microinjections, zygotes were washed 4–5 times in KSOM and placed back into culture. H3.3-GFP fluorescence was verified at the 2-cell stage as a marker for injection.

Given an estimated microinjection volume of 10 picoliter using the Femtojet parameters described above, the epididymosomal small RNA concentration of 1 ng/μl was chosen to achieve delivery of roughly a single sperm’s payload of small RNAs to a given zygote. Unpublished titrations in parthenote injections suggest that small RNA effects on Caput-specific genes are robust over at least a 4–fold concentration range.

### Single embryo RNA sequencing

Single embryo RNA-Seq was carried out using the SMART–Seq protocol (Ramskold et al. 2012). Data were mapped using RSEM after removing microRNAs, snoRNA, rRNA, and PCR duplicates. Embryos with fewer than 10,000 detected transcripts were removed from the dataset. For most analyses we focused on mRNAs with a median abundance of at least 10 tpm across the dataset.

### HNRNPAB Immunofluorescence

Caput and cauda ICSI embryos were collected at the mid-blastocyst stage and fixed in 4% paraformaldehyde. Immunofluorescence staining was performed as described in (Torres–Padilla et al. 2006). The primary antibody used was Alexa Fluor–647 conjugated anti–HNRNPAB (ab210053).

### E7.5 Embryo Dissection

Noon of the day that a mating plug was observed was considered embryonic day 0.5 (E0.5) of gestation. E7.5 embryos were dissected in DMEM (Dulbecco’s Modified Eagle Medium) (Invitrogen, Cat. No. 31600-034) containing 10% heat inactivated fetal bovine serum (Atlanta Biologicals, Cat. No. S11150), penicillin (100 U/ml), streptomycin (100 μg/ml) (Invitrogen, Cat. No. 15140-122) and 20 mM HEPES (Fisher, Cat. No. BP310).

## SUPPLEMENTAL MATERIALS

### SUPPLEMENTAL FIGURE LEGENDS

**Supplemental Figure S1.**
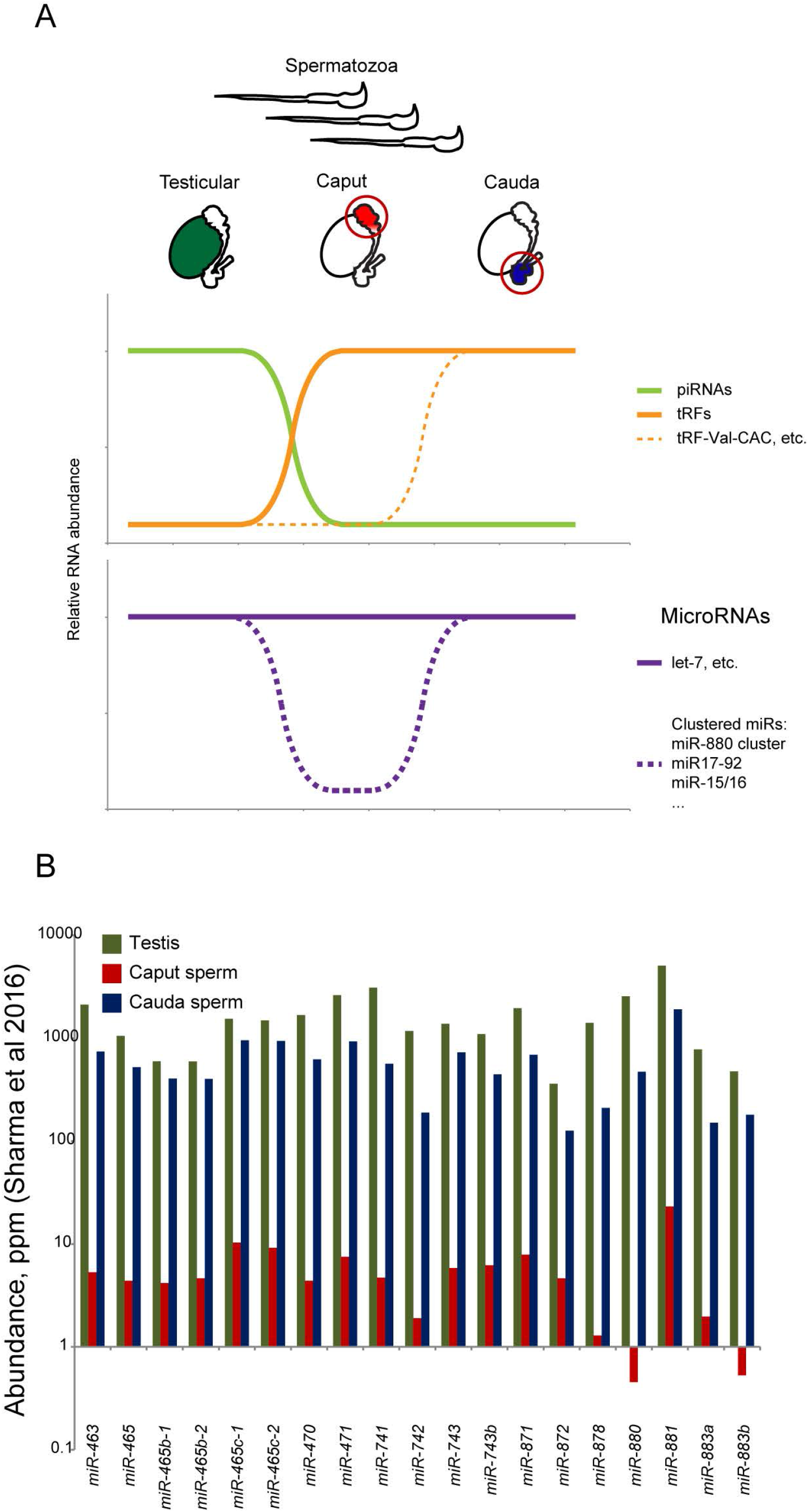
Schematic of small RNA changes during sperm development and maturation. A) Top cartoon shows three populations of spermatozoa, from the testis, caput epididymis, and cauda epididymis. Bottom panels show schematics of the major small RNA changes across this developmental trajectory, such as the global transition from piRNAs to tRFs occurring as sperm enter the epididymis. Note that while the majority of tRFs have been shipped to sperm already in the caput epididymis, several highly abundant tRFs (including tRF-Val-CAC, tRF-Val-AAC, and others – (Sharma et al. 2016)) are cauda-specific. Bottom panel shows schematic for microRNA dynamics in the epididymis. While many singleton microRNAs are relatively stable in abundance across these sperm populations, caput sperm do lose a wide variety of microRNAs (Nixon et al. 2015; Sharma et al. 2016), almost all of which are encoded in genomic clusters ranging from clusters of two (miR-15/16, etc.) to 100 kb-scale clusters with close to 20 miRNAs, such as the imprinted X-linked miR-880 cluster. Sperm transiting the epididymis then apparently re-gain a second round of these clustered microRNAs, which occurs via trafficking of epididymosomes from the surrounding epididymal epithelium (Sharma et al. 2016). B) Example of microRNA cluster dynamics in the epididymis. Data from Sharma *et al* 2016 show abundance (in parts per million reads, on a log_10_ y axis) for microRNAs comprising the miR-880 cluster in testis, caput sperm, and cauda sperm. Caput to cauda differences in these and other clustered microRNAs have also been documented in (Nixon et al. 2015), and we have reproduced these observations and confirmed that these microRNAs are highly expressed in purified testicular germ cell populations including mature testicular spermatozoa (see accompanying manuscript).

**Supplemental Figure S2.**
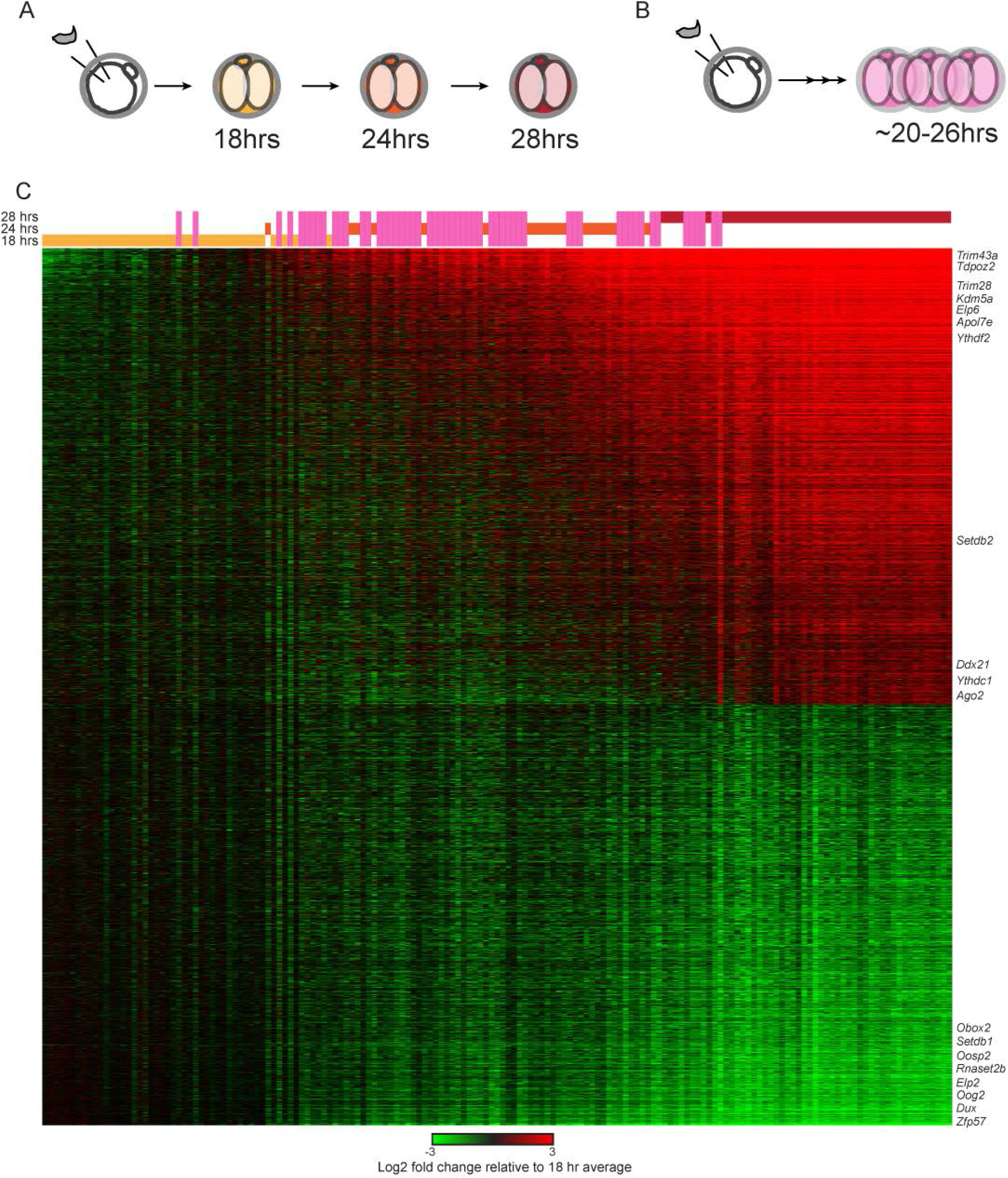
Zygotic genome activation. A) Embryos were generated using ICSI and cultured for precisely 18, 24, or 28 hours following sperm injection into the oocyte. Embryos were then promptly harvested for single-embryo RNA-Seq. B) In an original pilot experiment, ICSI embryos were generated over the course of roughly six hours on a given day (on several different days), then harvested for RNA-Seq 20 hours after the last sperm injection. These embryos thus span a poorly-staged time window of roughly 20-26 hours post-fertilization, providing samples that “fill in” temporal gaps between the 18, 24, and 28 hour embryos from (A). C) All 2-cell stage RNA-Seq. Embryos (columns) are sorted according to their correlation with the final ZGA (28 hours/18 hours) profile. The time of embryo collection is indicated as bars above the heatmap – long pink bars correspond to the relatively unstaged embryos.

**Supplemental Figure S3.**
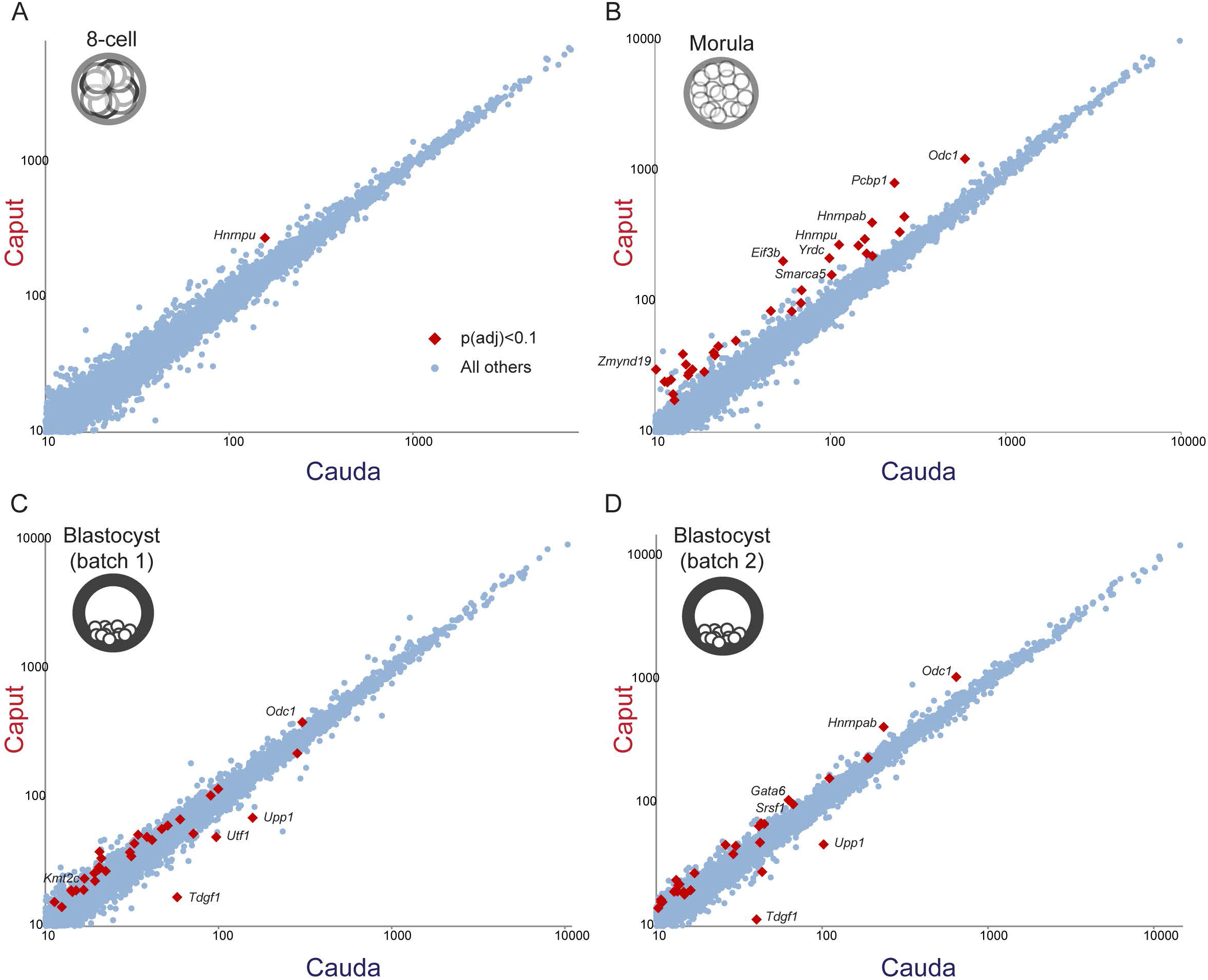
Caput vs. Cauda comparisons for 8-cell, morula, and blastocyst stages. Scatterplots as in **Figure 2A**, for the indicated developmental stages. Two separate biological replicates, carried out nearly one year apart, are analyzed separately for the blastocyst stage.

**Supplemental Figure S4.**
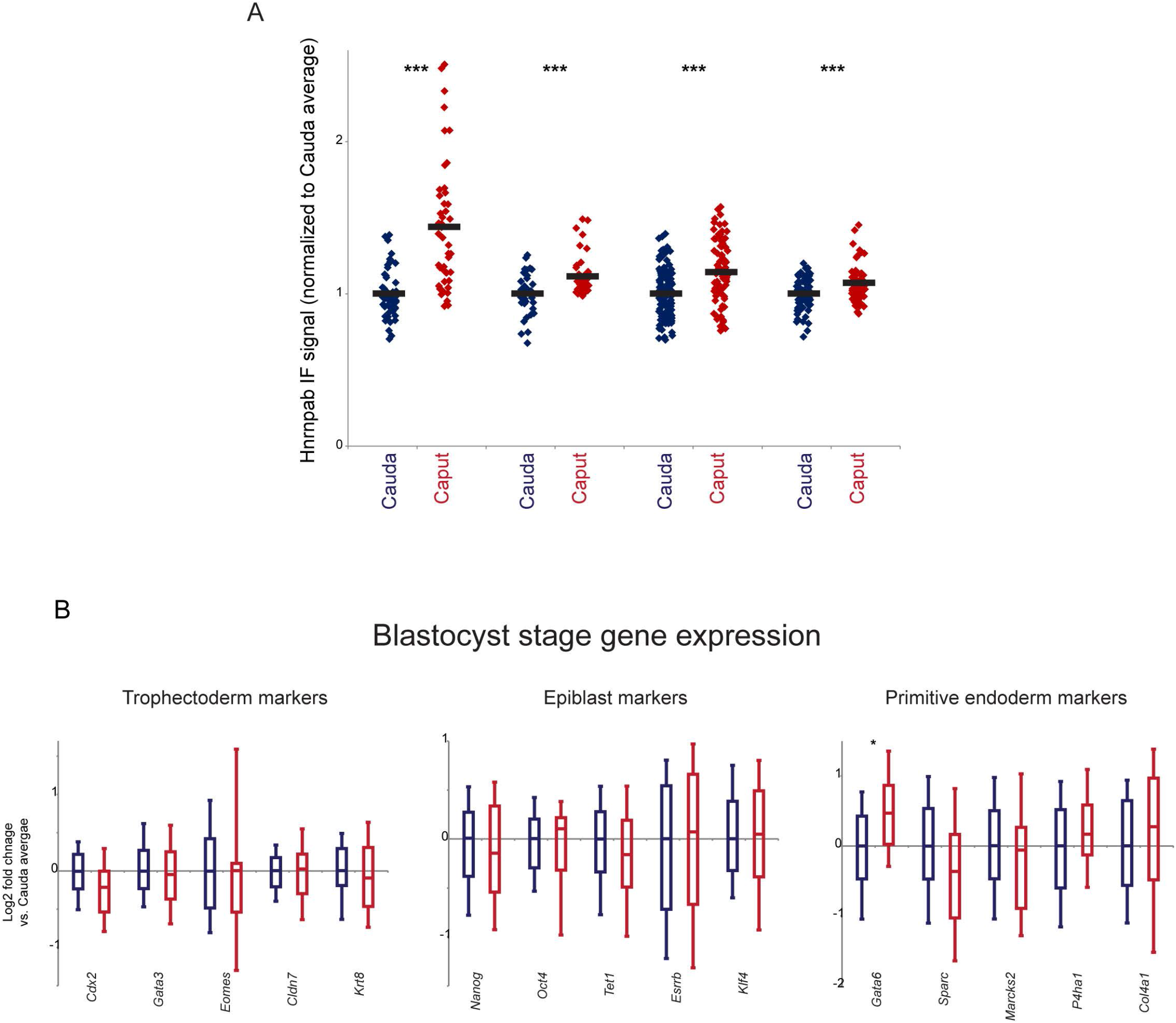
Validation and further analysis of Caput embryo regulatory phenotypes. A) Immunofluorescence (IF) validation of Caput effects on HNRNPAB protein expression in the blastocyst. Four panels show HNRNPAB levels, normalized to the Cauda average, for ~50-150 individual nuclei (from at least eight blastocysts for each experiment) for cauda- and caput-derived blastocysts. B) Sperm origin does not affect cell fate allocation in blastocysts. Box plots are shown as in **Figure 2C** for three sets of lineage marker genes in blastocyst RNA-Seq data, as indicated. With the exception of the significant upregulation observed for the gene encoding the key primitive endoderm transcription factor GATA6, we find no evidence for changes in cell composition between Caput and Cauda blastocysts.

**Supplemental Figure S5.**
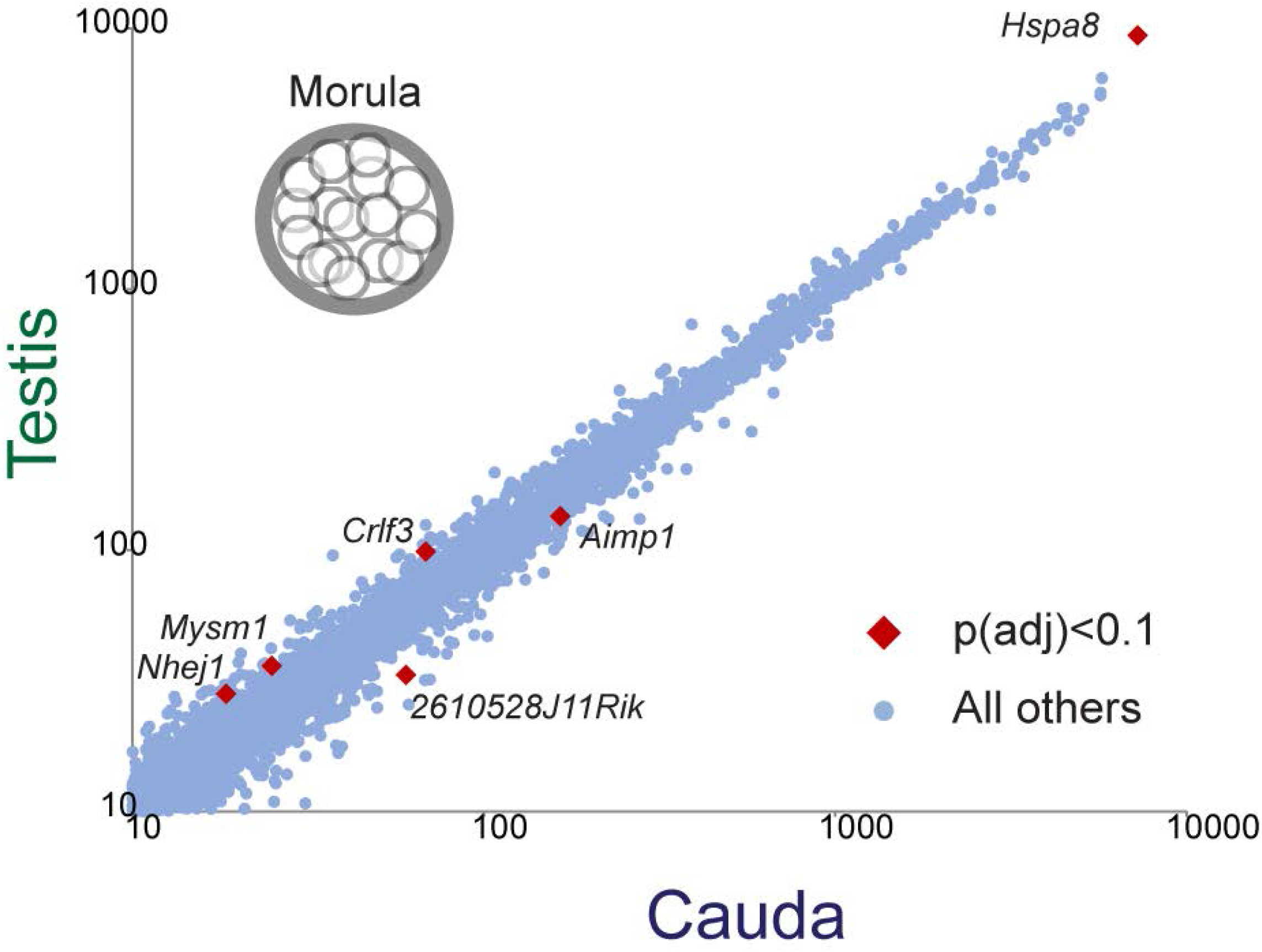
Morula-stage gene expression in testicular ICSI embryos. Scatterplot comparing median mRNA abundance in Cauda (x axis) and Testicular ICSI (y axis) embryos, as in **Figures 4C-D**. More detailed analysis of this experiment will be presented elsewhere – most relevant for this study is the absence of the Caput-specific regulatory program in embryos generated using testicular spermatozoa.

**Supplemental Figure S6.**
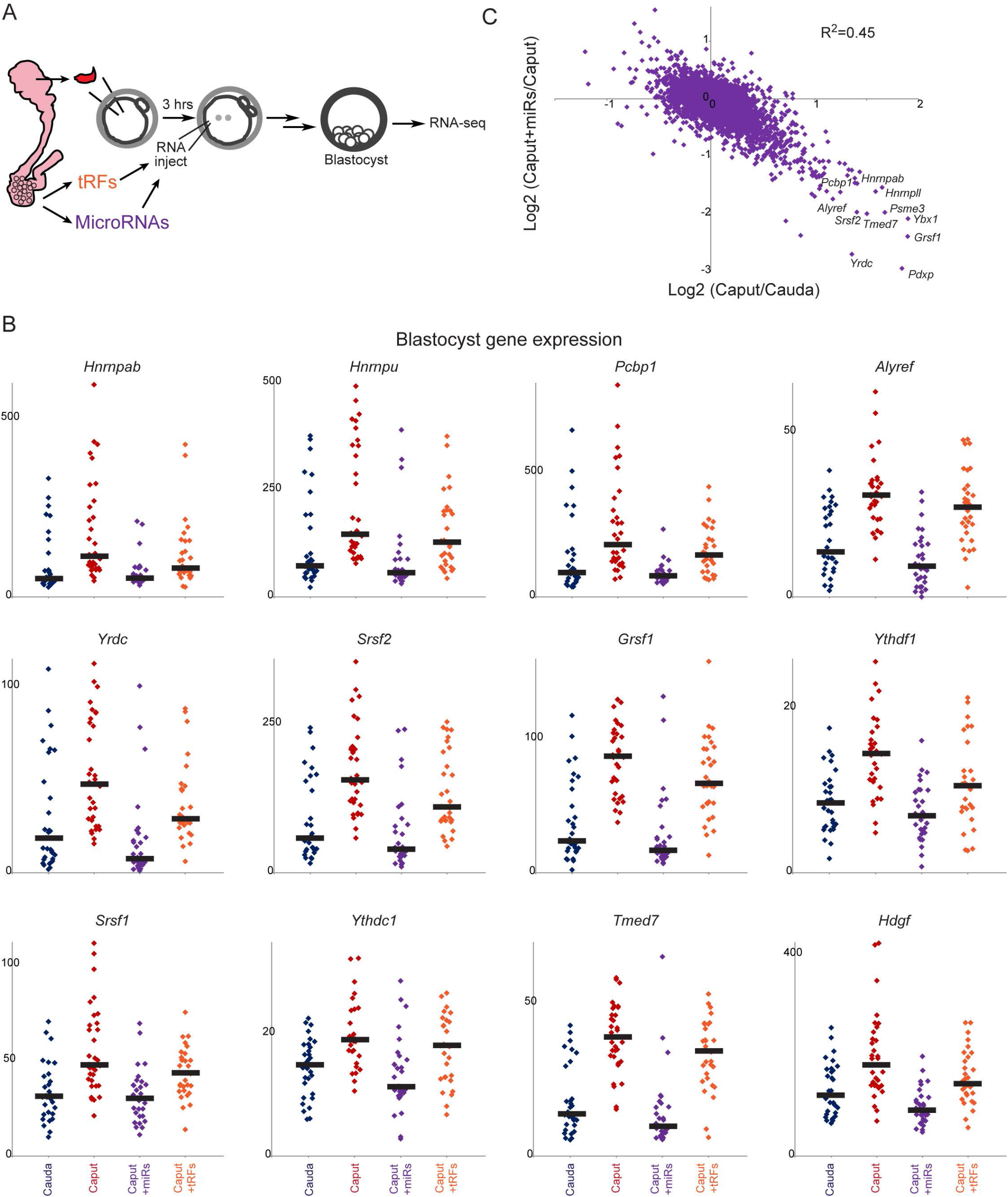
Cauda-specific microRNAs rescue Caput defects in blastocyst-stage gene regulation. A) Schematic of microinjection experiments. As in **Figure 6A**, but embryos were cultured to the blastocyst stage for single-embryo RNA-Seq. B) Data for individual blastocyst-stage embryos, plotted as in **Figure 6C**. C) Scatterplot comparing Caput effects on mRNA abundance (x axis) with effects of cauda-specific microRNAs (y axis), plotted as in **Figure 6D**.

### SUPPLEMENTAL TABLES

**Table S1. Fertilization and blastocyst progression**.

**Table S2. 2-cell stage gene expression**. Normalized single embryo RNA-Seq dataset for all 2-cell stage embryos.

**Table S3. 4-cell through morula stage gene expression**. Normalized single embryo RNA-Seq dataset for all 4-cell, 8-cell, and morula stage embryos.

**Table S4. Blastocyst stage gene expression**. Normalized single embryo RNA-Seq dataset for all blastocyst stage embryos.

**Table S5. Gene expression in small RNA-microinjected ICSI embryos**. Normalized single embryo RNA-Seq dataset for blastocysts, with Cauda, control Caput, and Caput+epididymosomal RNA embryos.

**Table S6. Gene expression in microRNA/tRF-microinjected ICSI 4-cell embryos**. Normalized single embryo RNA-Seq dataset for 4-cell stage embryos, with Cauda, control Caput, Caput+microRNA, and Caput+tRF embryos.

**Table S7. Gene expression in microRNA/tRF-microinjected ICSI blastocysts**. Normalized single embryo RNA-Seq dataset for 4-cell stage embryos, with Cauda, control Caput, Caput+microRNA, and Caput+tRF embryos.

**Table S8. Embryo numbers for all experiments**. Numbers of embryos used in the various RNA-Seq and embryo transfer experiments throughout this study.

